# The Neuroscience Multi-Omic Archive: A BRAIN Initiative resource for single-cell transcriptomic and epigenomic data from the mammalian brain

**DOI:** 10.1101/2022.09.08.505285

**Authors:** Seth A. Ament, Ricky S. Adkins, Robert Carter, Elena Chrysostomou, Carlo Colantuoni, Jonathan Crabtree, Heather H. Creasy, Kylee Degatano, Victor Felix, Peter Gandt, Gwenn A. Garden, Michelle Giglio, Brian R. Herb, Farzaneh Khajouei, Elizabeth Kiernan, Carrie McCracken, Kennedy McDaniel, Suvarna Nadendla, Lance Nickel, Dustin Olley, Joshua Orvis, Joseph P. Receveur, Mike Schor, Timothy L. Tickle, Jessica Way, Ronna Hertzano, Anup A. Mahurkar, Owen R White

## Abstract

Scalable technologies to sequence the transcriptomes and epigenomes of single cells are transforming our understanding of cell types and cell states. The Brain Research through Advancing Innovative Neurotechnologies (BRAIN) Initiative Cell Census Network (BICCN) is applying these technologies at unprecedented scale to map the cell types in the mammalian brain. In an effort to increase data FAIRness (Findable, Accessible, Interoperable, Reusable), the NIH has established repositories to make data generated by the BICCN and related BRAIN Initiative projects accessible to the broader research community. Here, we describe the Neuroscience Multi-Omic Archive (NeMO Archive; nemoarchive.org), which serves as the primary repository for genomics data from the BRAIN Initiative. Working closely with other BRAIN Initiative researchers, we have organized these data into a continually expanding, curated repository, which contains transcriptomic and epigenomic data from over 50 million brain cells, including single-cell genomic data from all of the major regions of the adult and prenatal human and mouse brains, as well as substantial single-cell genomic data from non-human primates. We make available several tools for accessing these data, including a searchable web portal, a cloud-computing interface for large-scale data processing (implemented on Terra, terra.bio), and a visualization and analysis platform, NeMO Analytics (nemoanalytics.org).

**KEY POINTS:**

- The Neuroscience Multi-Omic Archive serves as the genomics data repository for the BRAIN Initiative.
- Genomic data from >50 million cells span all the major regions of the brains of humans and mice.
- We provide a searchable web portal, a cloud-computing interface, and a data visualization platform.

## INTRODUCTION

Circuits in the mammalian brain are comprised of billions of neurons, connected via trillions of synapses. Neuroscientists have long recognized that the structural and functional properties of brain circuits arise, in part, from the diverse anatomical, physiological, and molecular characteristics of their composite neuronal and non-neuronal cells. Surprisingly, however, precise definitions for the myriad subtypes of brain cells have remained elusive (1, 2).

Single-cell genomics has emerged as a powerful and scalable technology to more rigorously define the brain’s cell types (3–5). Single-cell and single-nucleus RNA sequencing (scRNA-seq) have been applied to sequence the transcriptomes of millions of cells in mammalian brain regions, identifying hundreds or even thousands of transcriptionally distinct cell types (1–6). Multimodal technologies to co-assay a cell’s transcriptome along with its morphology, physiological characteristics, or spatial location have made it possible to relate these transcriptomic cell types to classically defined anatomical and functional cell types (7, 8). Single-cell epigenomic technologies characterize the cell type-specific gene regulatory mechanisms underlying cell type identity (9, 10).

The Brain Research through Advancing Innovative Neurotechnologies (BRAIN) Initiative is applying single-cell multi-omic techniques at unprecedented scale to map the cell types in the mammalian brain (11, 12). Since 2017, the BRAIN Initiative Cell Census Network (BICCN) has worked to generate an open-access reference atlas, integrating molecular, spatial, morphological, connectomic, and functional data to describe the cell types in mouse, human and non-human primate brains (11). In its first phase, the BICCN produced detailed atlases for the cell types in the primary motor cortex (12–14), as well as for the cortex’s prenatal development (15, 16). In its continuing work, the BICCN is nearing the completion of draft cell type atlases for nearly all brain regions in mice and humans. These efforts are expected to be refined over the next several years and expanded to additional conditions through BICCN’s successor, the BRAIN Initiative Cell Atlas Network.

An integral goal of the BICCN is to make these brain cell resources available for use by the broader research community, consistent with data FAIRness (Findable, Accessible, Interoperable, Reusable) (17). Toward this goal, the National Institutes of Health have funded the creation of data archives and tools for the web-based analysis and visualization of BRAIN Initiative data. Here, we describe the Neuroscience Multi-Omic Archive (NeMO Archive), which serves as the primary genomic data repository for the BRAIN Initiative. The NeMO Archive provides access to an expansive collection of single-cell genomic data from the mammalian brain, including a variety of tools for searching, downloading, and analyzing these data. A companion website, NeMO Analytics, provides additional, biologist-friendly tools for analysis and visualization.

## MATERIALS AND METHODS

### Data deposition and data processing backend

The NeMO Archive receives submissions of genomic data from BRAIN Initiative researchers, as well as other authorized contributors. Data submission begins with the upload of a standardized manifest file that submitters create based on a provided template. The manifest outlines all files that will be part of the submission and includes required metadata. Once the manifest is validated for completeness and adherence to controlled vocabularies, submitters receive a directory name and command with which to submit data directly to the NeMO Archive using the IBM Aspera data transfer tool. NeMO servers contain three “incoming” or landing areas, the public incoming area for data with no restrictions that are slated for immediate public release, the restricted incoming area for data requiring consented access (including controlled access human data), and the embargo incoming area for data to be held in embargo before publication. Submitters notify NeMO that their upload has completed through the submission of a tagged file, at which point automated processes pick up the submission for initial quality assurance, including detection of all expected files and checksum comparisons to ensure that files were not corrupted during data transfer. We note that no BICCN data are subject to an embargo.

### Metadata

Data must be presented together with detailed metadata in order to be useful to the wider neuroscience research community (17). Toward this end we have worked with the BICCN infrastructure working group and the larger neuroscience community to ensure that we capture all essential and relevant metadata in the archive, including information about the data source, organism, the experimental assays used, and the sequencing technologies and bioinformatics tools used to generate the data and derived results. We are working to implement the use of standard ontologies and controlled vocabularies including NCBI Taxonomy (18), OBI (19), EDAM (20), Uberon (21) to capture information about organism, experimental assays, file types and formats, and anatomy respectively.

### Identifiers, Datasets and Releases

An important element of the FAIR principles is to ensure that data are findable, both for human operators as well as algorithms. We assign a stable local unique identifier to each subject, sample, and file asset at NeMO. This ensures that we can track each of these assets uniquely over the lifetime of the asset. We have registered NeMO as a source of such identifiers with identifiers.org to ensure NeMO identifiers can be resolved by the identifier resolver service. In addition to assigning local identifiers to individual assets, we also assign local as well as global unique identifiers to asset collections, which might include data used in a paper or a NeMO quarterly data release. These asset collections are released as BDBags (22), a structured method to create a collection of objects, and each of these is assigned a digital object identifier (DOI) issued through DataCite (23). FAIR principles require that an identifier can be used to find additional information about the object that can be consumed by both people and machines. In this spirit, each NeMO asset has a landing page which describes the asset for humans, as well as a structured JSON object describing the asset for machines.

### Consensus data processing

Consensus processing of BICCN data was performed using workflows implemented on Terra, a cloud-native workbench developed by Broad, Verily, and Microsoft (Terra.bio). These data include single cell transcriptomic data generated using 10× Genomics and SMART-seq technologies, single cell methylation data (snmC-seq), and single cell chromatin accessibility data (snATAC-seq). Single-cell and single-nucleus 10x Genomics transcriptomic read-level data were processed to generate exon/intron counts using the Optimus pipeline (24) (RRID:SCR_018908). SMART-seq data were processed with the Smart-seq2 Single Nucleus Multi-Sample Pipeline (25) (RRID:SCR_021312) in batches of 100, optimized to maximize the number of samples run on a single virtual machine within 24 hours and minimize cost. For consistency, the same genome build and gene annotation were used for 10X and SMART-seq processing. snATAC-seq reads were aligned using the scATAC Pipeline (26) (RRID:SCR_018919), which utilizes SNAP-ATAC (27) and summarizes read counts per 5000 base pairs. snmC-seq data were processed using the CEMBA pipeline (28, 29) (RRID:SCR_021219), which utilizes bowtie2 for alignment and produces counts for every CpG site in the genome.

### NeMO Portal

To facilitate the exploration of data acquired by the NeMO Archive, the project has provisioned a web application called the NeMO Portal, available at https://portal.nemoarchive.org. The portal allows users to explore the data in various usage patterns. For example, the portal allows users to filter the existing data using facets along the left side of the interface. This is known as “faceted search” and is commonly used in well-known e-commerce websites such as amazon.com. By using faceted search, we present a familiar and accepted usage pattern to end users of the portal to allow them to intuitively narrow down the available data to only the datasets and files that they are interested in. Facets include organism (e.g., mouse, human, marmoset), brain region, sequencing technique, and modality type (transcriptomics, epigenomics), among others. The site is dynamic in nature such that when different facets are selected or de-selected, updated charts and visualizations about the filtered data are automatically re-generated and presented to the end users so that they may better comprehend how the facets affect their selections.

The portal also provides an “Advanced Search” interface, which allows users to directly enter a query using a simple query language. If a syntax error is detected, the portal’s interface reports where in the query the error is located for easier correction.

Charts and graphs update dynamically as the query is refined. The use of the aforementioned facets is also translated into an advanced search query statement.

When users have completed their exploration of the data and found datasets or files they are interested in, the portal allows them to add them (individually or in bulk) to a data cart. Again, the usage is similar to common patterns seen elsewhere on the web for greater familiarity. The cart has features that allow the user to download a file, called a manifest, which contains metadata and the network locations of the files of interest. To download these files, additional ancillary tools associated with the portal have been developed, such as the portal client (https://github.com/IGS/portal_client), that allow the files in the manifests to be downloaded. This approach is taken because it is not feasible to dynamically download the user’s selections directly to the browser, as the total volume can easily become tens of terabytes of size. The NeMO portal has also been integrated with the Terra platform to allow manifests to be exported into a Terra workspace for processing on the cloud, avoiding the need for local data download.

### NeMO Analytics

The NeMO Analytics portal is implemented as an instance of the Gene Expression Analysis Resource (gEAR) software (30) for visualization and analysis of transcriptomic and epigenomic data. A strength of the portal is in its ability to display multiple multi-omic datasets in a single page, effectively allowing a page (termed ‘profile’) to fully support a manuscript or include a collection of thematically related datasets. Direct access to the data is often offered by custom URLs presented in manuscript figure legends and text. Finally, data analysis tools are attached to each dataset, allowing further exploration of the data also to biologists with limited informatics skills (31). Data from BICCN were obtained from data generating labs via the NeMO Archive. Non-BICCN data were obtained from public repositories, including the Gene Expression Omnibus, the UCSC Cell Browser (cells.ucsc.edu), and GEMMA (32). Data were uploaded into the portal via the Data Uploader. For sn/scRNA-seq, Patch-seq, and MERFISH experiments, we uploaded read counts and associated metadata. For scATAC-seq and scMethyl-seq experiments, we created TrackHubs and linked them to the portal. Custom visualizations for each dataset were created using the Data Curator tool and assembled into Profiles, enabling several datasets to be viewed side-by-side.

## RESULTS

### A resource of single-cell multi-omic data from the brains of humans, non-human primates, and mice

The NeMO Archive has been continuously ingesting single-cell transcriptomic and epigenomic data from BICCN researchers since August 2018. As of August 10, 2022, the archive contains 383.1 terabytes of data in 1,216,873 files from 289,707 biological samples (Fig. 1A). In total, these data describe the transcriptomes or epigenomes of approximately 57,452,822 cells.

**Figure 1.**
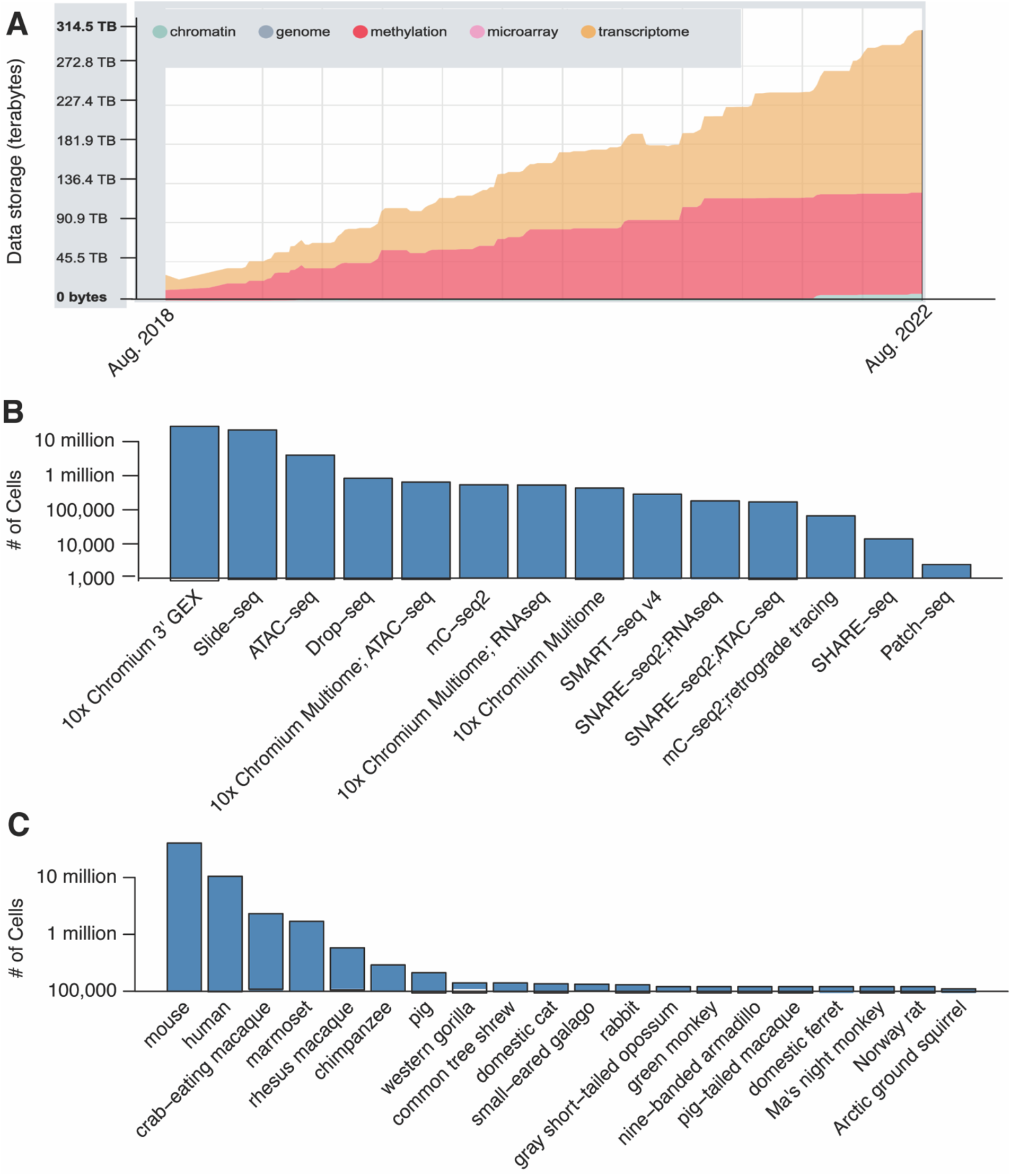
NeMO Archive data inventory. A. Growth of data storage in the NeMO Archive over time. B. Counts of cells assayed by each transcriptomic or epigenomic technology. C. Counts of cells for each species. Y-axes in panels B and C are on a log-scale.

BRAIN Initiative researchers utilized several distinct single-cell technologies to profile brain cell types (Fig. 1B). Single-cell transcriptomic data (scRNA-seq) were generated from 27,921,806 cells/nuclei with variations of the 10x Genomics 3’ Gene Expression technology (33), which captures cells via droplet microfluidics and generates sequence from the 3’ ends of RNA molecules. scRNA-seq was generated from an additional 834,367 cells with Drop-Seq, which is also a droplet-based method (3). Fulllength transcriptomes were generated from 286,418 cells and nuclei with SMART-seq (34). Spatial transcriptomic data were generated from 21,802,314 spatial positions in the brain at 10 um resolution using Slide-seqV2 (35) (henceforth, spatial positions are included among summaries of cell counts as single-cells, consistent with the near-single cell resolution of these data). Single-cell epigenomic data include open chromatin profiling of 4,021,312 cells with the Assay for Transposase-Accessible Chromatin (scATAC-seq) (36), as well as single-cell profiling of DNA cytosine methylation (snmC-seq2) (37) in 539,969 cells. Single-cell ATAC and RNA multi-omes were generated from 1,161,948 cells using 10x Genomics Multi-ome, SHARE-seq (38), and SNARE-seq (39). A subset of the SMART-seq and snMethyl-seq data were generated as part of multimodal experiments, including imaging of neuronal morphology and projection patterns, retrograde tracing, and electrophysiology (Patch-seq). Summaries of all the BICCN projects contributing data to the NeMO Archive are available at biccn.org.

Data in the NeMO Archive were derived from the brains of 20 mammalian species (Fig. 1C). The majority of the data are from laboratory mouse strains (40,232,712 cells). These are primarily from 8-week-old mice of the C57BL/6 strain and its derivatives expressing cell type-specific Cre reporters, and a smaller number of samples describe pre- and postnatal mouse brain development. Samples from mice are annotated to 194 distinct sub-regions and cell populations labeled by Cre reporters, spanning all of the major structures of the forebrain, midbrain, and hindbrain, as defined in the Allen mouse brain common coordinate framework (40). 10,503,719 cells are from the human brain. These data include extensive atlases for both the adult and developing human brain and are annotated to 269 distinct sub-regions and cell populations. Adult samples were derived from donors who were 18-68 years old at time of death with no history of psychiatric or neurological disorders. Developmental samples span the full range of human prenatal development from 4-34 gestational weeks, as well as early postnatal development. The BICCN has also generated extensive single-cell genomic resources for the brains of non-human primates and other mammals. These include atlases for many of the brain regions of marmosets (1,693,284 cells) and rhesus macaques (579,187 cells), as well as surveys of specific forebrain structures in 16 additional species spanning several mammalian clades.

While a subset of these data have been presented in peer-reviewed publications, the majority of the data are being released to the research community prior to publication by the BICCN consortium in an effort to accelerate discovery. Data producers within BICCN have submitted, at a minimum, raw data consisting of FASTQ files, as well as metadata describing the species, genotype, brain region, technique, investigator, grant number, and related information about each sample. The NeMO Archive also provides access to processed data such as read counts and cell type assignments from BICCN publications.

Each sample and file in the NeMO Archive is assigned a unique identifier and indexed by its metadata (Methods). We provide multiple interfaces to the data (Fig. 2), including direct access to the data via http, sftp, and Google Cloud Platform buckets; a web portal enabling users to search for datasets and download them (nemoarchive.org), a cloud-computing interface enabling users to perform data processing at scale (using pipelines implemented in Terra, terra.bio), and web-based visualization and analysis tools implemented at a companion website, NeMO Analytics (nemoanalytics.org). These interfaces are described in detail below.

**Figure 2.**
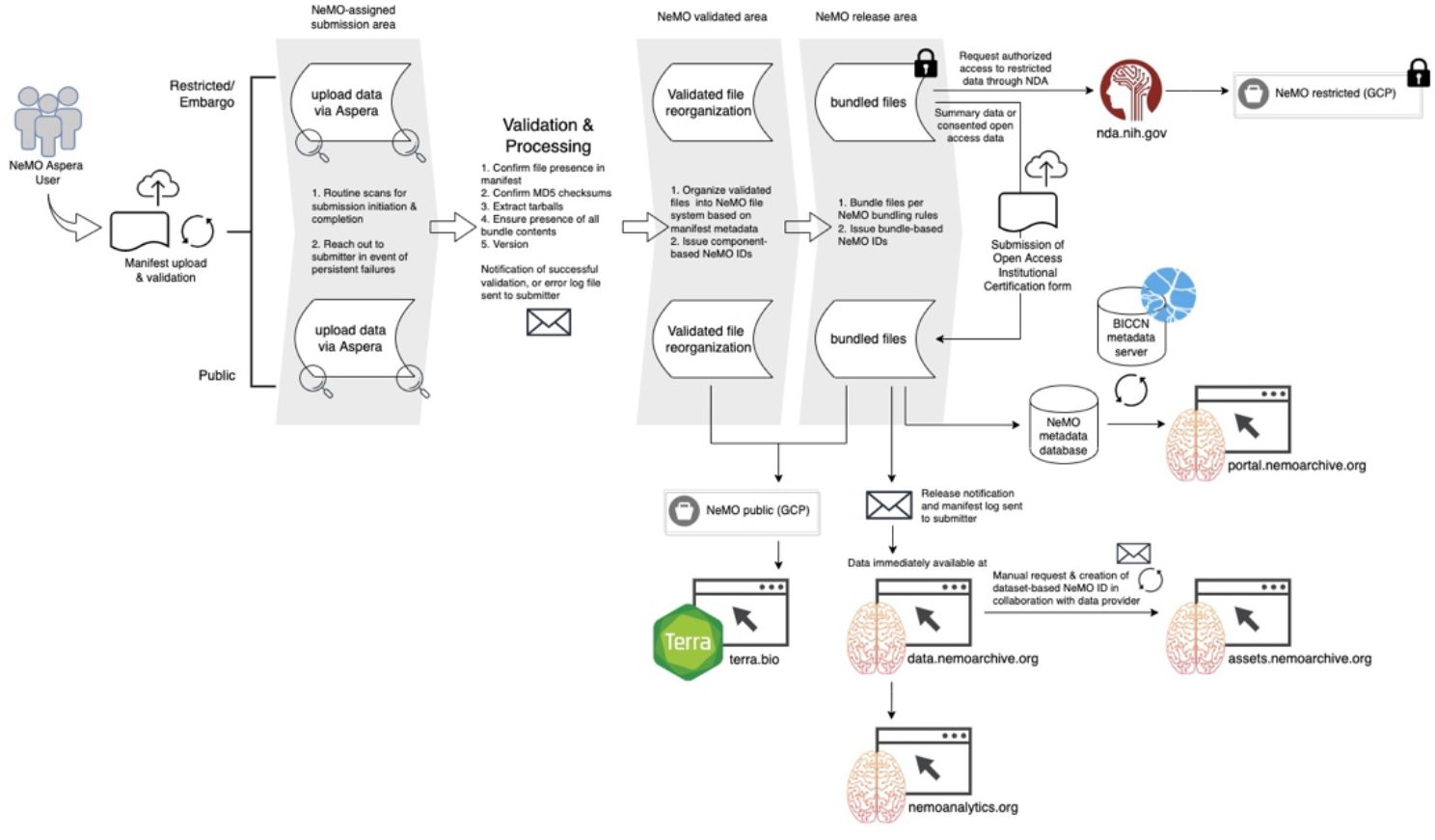
Data processing workflow. Flow of data from ingest through validation and release to NeMO resources.

### Consensus processing of multi-omic data with the BICCN cloud-computing environment

To facilitate joint analyses of BICCN data, we have developed consensus pipelines for large-scale processing of single-cell genomic data submitted to the NeMO Archive. These pipelines are implemented on Terra in partnership with BICCN’s Analysis Working Group and written in Workflow Description Language (WDL), a workflow processing language designed to be easy for humans to read and write. WDL provides the flexibility to combine multiple computing languages in a single workflow and execute this workflow on a variety of local or cloud platforms. Several BICCN pipelines are available, including Optimus, a pipeline for processing 10x Genomics scRNA-seq and snRNA-seq data; a full transcript (SMART-seq) scRNA-seq and snRNA-seq data processing pipeline (Smart-seq pipeline); and a pipeline for snATAC-seq utilizing SNAP-ATAC (27). Each cloud-native pipeline was rigorously tested by consortium members to ensure outputs replicated expert in-house pipelines and optimized to improve performance and cost in the cloud environment. For example, the cost of SMART-seq processing was decreased 50-fold by switching from HISAT2 (41) to STAR (42) for genome alignment, using dedicated virtual machines with pre-loaded reference genomes, and optimizing the number of samples to process within a 24-hour period to maximize compute resources. Additionally, engineers from Terra and NeMO utilized the APIs to submit hundreds of jobs to Terra via the command line, facilitating large-scale processing and removing the need to use manual user interfaces. Each of these pipelines is publicly available on Terra and listed in the BICCN portal (https://biccn.org/tools/biccn-pipelines).

As of July 31, 2022, we have completed consensus processing using these pipelines for the adult mouse brain atlas, consisting of 1,475 10x Genomics sc/snRNAseq samples (~13,000,000 cells); 213,410 SMART-seq samples (213,410 cells); 138 single-nucleus methyl-cytosine sequencing (snmC-seq) and 185 epi-retro seq samples (168,098 and 19,357 cells respectfully); and 96 snATAC-seq samples (1,094,579 cells). In addition, we make available read counts from the adult and prenatal human brain atlases, processed through equivalent pipelines with CellRanger. All of these processed data are available in the NeMO portal (see below).

### Accessing BICCN data through the NeMO Portal

NeMO has deployed a data portal where users can search for and download data. The home page of the NeMO data portal (Fig. 3A) gives the data summary including the number of grants or studies represented at NeMO, the file count summarized by the data modality for each of the grants, and quick links to featured datasets. The home page also serves as the launchpad to access other functions of the portal including accessing additional information about each of the grants/studies and accessing the data search and discovery page.

**Figure 3.**
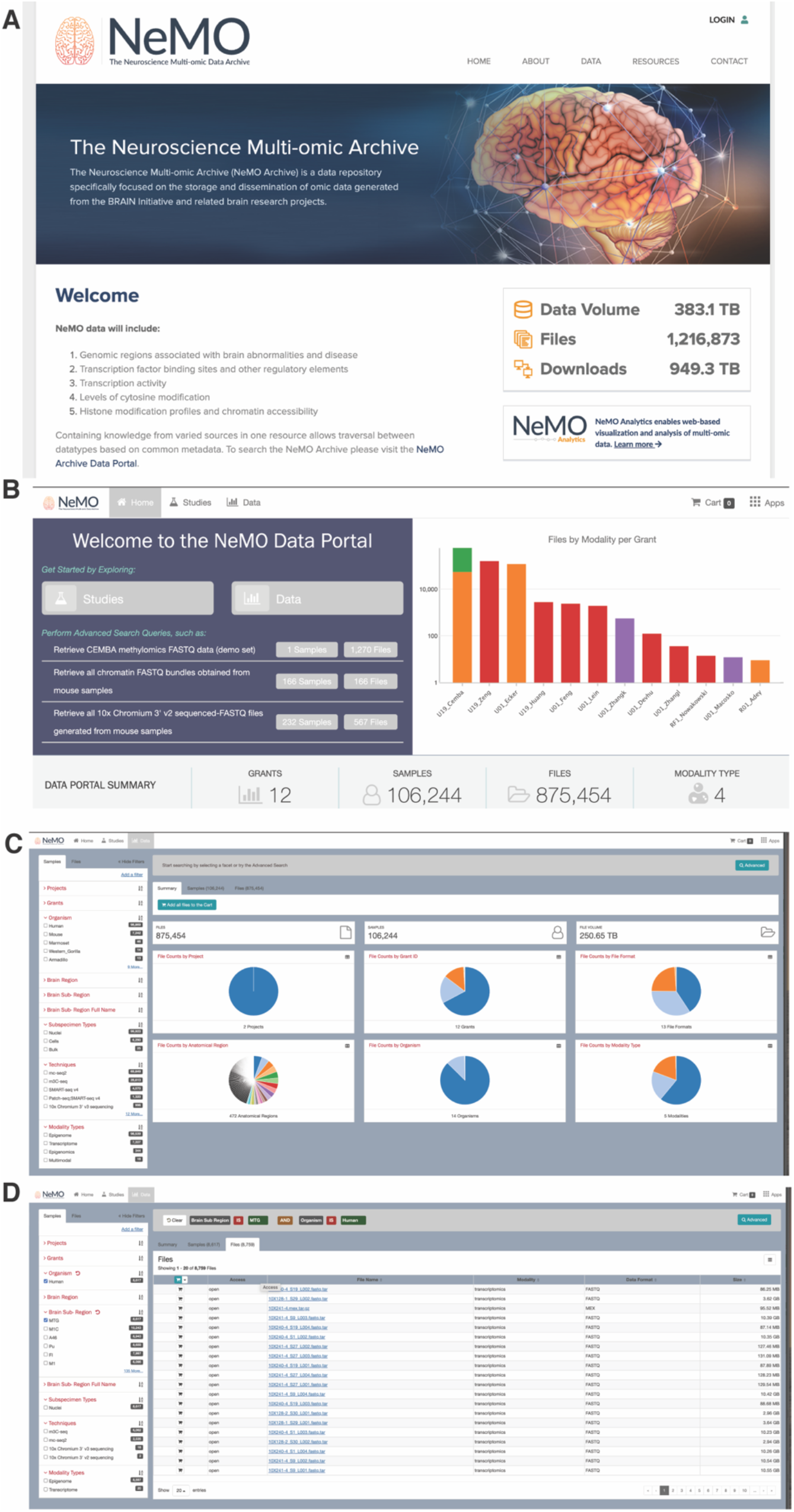
Accessing BICCN data through the NeMO Portal. A. Users can find and access data in the NeMO Archive through a website and portal (www.nemoarchive.org). B. The landing page allows users to get information about the project, look at summary of data available at the archive, growth of data over time at the archive, access documentation on tools and data downloads, and find links to other related resources. C. The data discovery and search page of the data portal provides a filtering interface (left) and summaries based on different metadata properties (right). Users use the faceted search interface to filter the data by selecting or deselecting the check boxes next to the facets. D. Once the user has selected a set of files or samples they can use the “shopping cart” features, accessed by clicking on the cart button on top right, for further actions.

The filtering interface organizes the facets in different categories and displays the items within a category. In addition, the interface displays the count of objects that are tagged with that facet. The filters can be applied to “Samples” or “Files” as represented by the two tabs in the top left corner. Sample-level filters are based on characteristics such as the project or study, the organism, the anatomical region, data modality, and the sequencing technique. File-level filters include characteristics such as file data type or file format. Besides the filters that are visible, the users can access additional filters by clicking on the “Add a filter” button to access other metadata fields that can be used to filter the data. As filters are applied, the summary information presented as charts on the right are automatically updated. In addition, the query that is being built through filtering is displayed in the query box on the top of the screen. The charts themselves are interactive and users can click on the charts to add additional filters. Users can view any of the charts as a table by clicking on the table icon on the top right of the chart panel. In addition to the faceted search, the portal also has an advanced search button that can be accessed by clicking on the “Advanced” button on the query box. This brings up the advanced search interface (See Fig. 3C) where users can use the type-ahead feature (a feature where users start typing a letter all possible fields starting with those letter(s) are displayed for users to choose) of the portal to filter data based on all the available metadata fields. For advanced users familiar with the metadata this can be a more efficient search interface. Another advantage of the advanced interface is that users can copy and save the text query search for subsequent use.

Once the user has narrowed down the dataset the user can reuse the “Samples” or “Files” tab (Fig. 3C) to select one or more of the objects to add to the “Data Cart”. If a sample is selected, all files associated with that sample are added to the data cart. If a file is selected, only the selected file is added to the cart. The data cart page gives a summary of the data including the number of files, samples associated with these files, the cumulative size of the data. In addition, the interface displays a list of the files in a table. Here the user can click on the hyperlink associated with a file to get additional information about the file (Fig. 3D) or use the “Download” button to either download the data or move the data to other linked resources such as the Terra cloud-computing environment. When the user chooses to download the data, the user must download a manifest file, which stores the information necessary to download the actual data to the local computer. A separate download tool is necessary to download the actual data. The user can also choose to download the metadata associated with the files in the data cart. If the user wishes to make use of the data with pipelines at Terra, no data is actually moved, rather only information necessary for Terra to access the data stored in the NeMO cloud is transferred to Terra.

### Visualization and analysis of multi-omic brain data with NeMO Analytics

It can be challenging for biologists not skilled in bioinformatics to fully utilize NeMO Archive data for visualization and analysis. To address this, we have initiated NeMO Analytics (nemoanalytics.org), a portal designed to allow a broad range of neuroscientists to fully benefit from these data without requiring any expertise in programming. NeMO Analytics gene expression tools are powered by gEAR (umgear.org) (30) and consist of the following six components: an Expression Browser allows the visualization of gene expression patterns (one gene at a time) across multiple datasets or views; the Curator Tool makes views of datasets customizable by the user, supporting data presentations as bar plots, line plots, x-y scatter plots (e.g., tSNE and UMAP plots of scRNA-seq data), violin plots, and SVG images; the Comparison Tool tests pairwise contrasts of all of the genes in two groups (e.g., two cell types); the Single Cell RNA-seq Workbench enables on-the-fly cell type clustering and marker gene detection via a Scanpy workflow (43); epigenomic data can be visualized in the context of a linear genome browser via integration with EpiViz (44); the Dataset Manager allows the selection of groups of datasets to visualize side-by-side.

We produced eight multi-dataset profiles enabling users to interact with the primary motor cortex datasets described in publications by the BICCN (12–14) (Fig. 4). The mouse transcriptomic, epigenomic, and “integrated” profiles provide visualizations of these data types from mouse primary motor cortex (14). The cross-species and merged-species profiles provide visualizations of transcriptomic data from integrated analyses of human, marmoset, and mouse primary motor cortex (13). The Patch-seq profile displays gene expression in the context of physiological cell types from multimodal transcriptomic and electrophysiological profiling of primary motor cortex neurons (12). The Spatial profile displays MERFISH single-cell resolution in situ transcriptomics data on cortical sections through the primary motor cortex (12). The Cortical Development profile displays single-cell and bulk tissue transcriptomic data from the developing human cortex at pre- and postnatal timepoints (16, 45–47).

**Figure 4.**
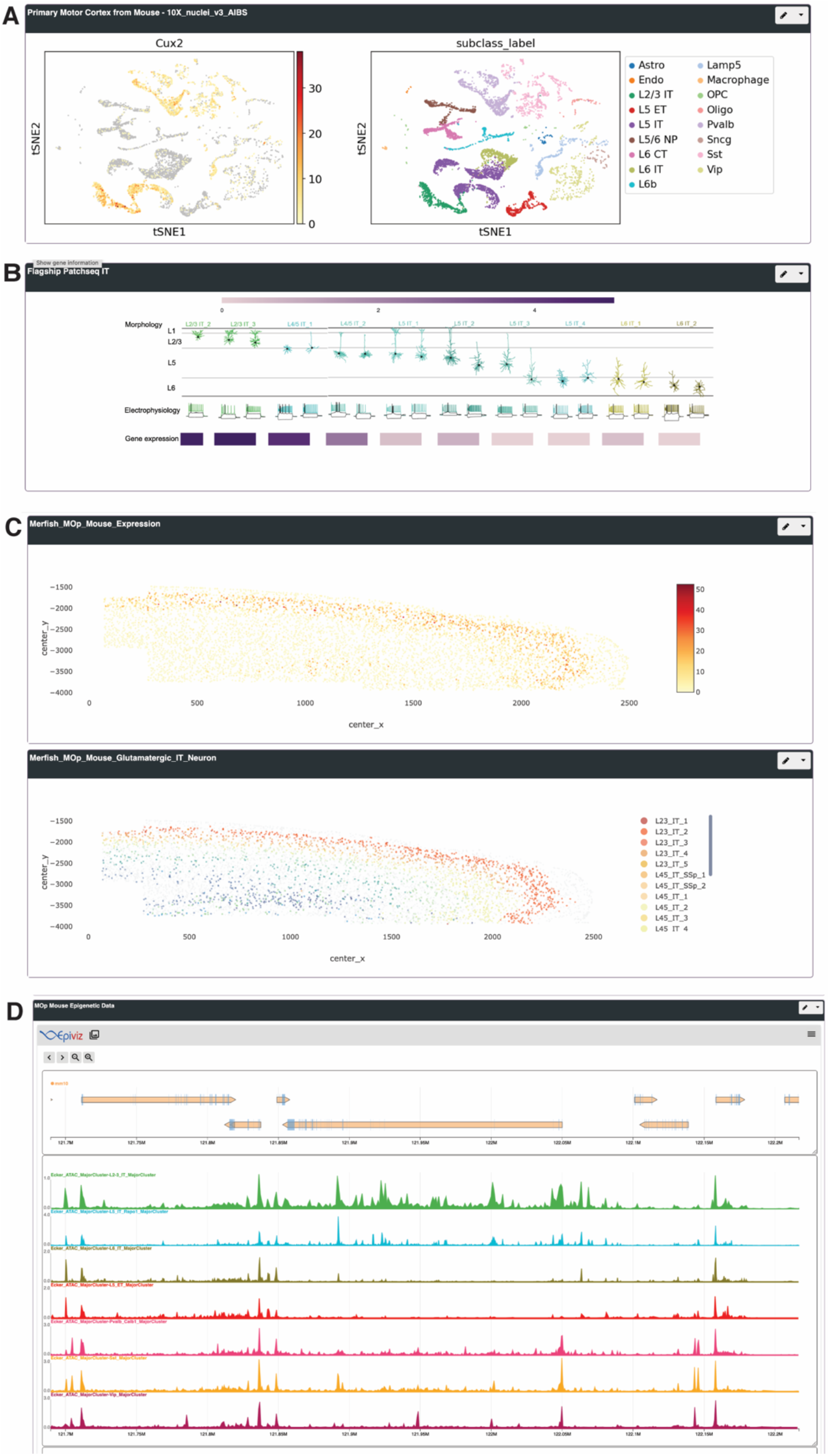
Visualization of BICCN data with NeMO Analytics. Expression of *Cux2,* a layer-specific marker of excitatory neurons, in BICCN data from the primary motor cortex, visualized with NeMO Analytics. A. clustering of transcriptomic cell types with 10x Genomics snRNA-seq (https://nemoanalytics.org/index.html?&layout_id=4e8f6c00&gene_symbol=Cux2). B. Multimodal characterization of gene expression, morphology, and electrophysiology with Patch-seq (https://nemoanalytics.org/index.html?&layout_id=1206f7ce&gene_symbol=Cux2). Spatial transcriptomics with MERFISH (https://nemoanalytics.org/index.html?&layout_id=7e14ad94&gene_symbol=Cux2). D. Chromatin accessibility profiling with scATAC-seq (https://nemoanalytics.org/index.html?&layout_id=2c72ff02&gene_symbol=Cux2).

A long-term goal for the BRAIN Initiative is to integrate cell type atlases with research on brain disorders. Toward this end, we have assembled several profiles with single-cell and traditional transcriptomics and epigenomics experiments related to Alzheimer’s disease (AD). These NeMO-AD profiles feature >50 AD-related RNA-seq and scRNA-seq datasets, including data from post-mortem brain tissue of AD cases vs. controls, as well as from animal models. Using these profiles, one can assess AD-related transcriptional changes across disease progression, in several brain regions, and in specific cell types. We have also created a NeMO-AD profile for the visualization of data describing microglial cell states, since AD is associated with brain inflammation mediated by these immune cells.

In addition to these pre-designed profiles, NeMO Analytics enables users to develop their own profiles from BICCN datasets in the NeMO Archive or by uploading any dataset of interest. Read counts from ~80 BICCN scRNA-seq experiments are available in the Single-Cell RNA-seq Workbench so that users can explore molecular cell types in individual samples. Users can then add their analyses to a profile or download the data to perform more detailed analysis. The Dataset Uploader allows users to create visualizations and analyses of any scRNA-seq, RNA-seq, or epigenomic dataset of interest. These datasets are initially private to the user. At their discretion, users can share their custom datasets and profiles with other users or make them fully public. The latter feature has proven useful in rapidly producing companion websites for both BICCN (12–14) and non-BICCN (48, 49) publications.

## DISCUSSION

The single-cell transcriptomic and epigenomic data generated by the BICCN and housed in the NeMO Archive are valuable resources to the research community. New features will be regularly added to the NeMO Portal and to NeMO Analytics with an eye toward maximizing the utility of these resources. We continue to ingest new data from BICCN and are expanding our work to incorporate data from other sources. Most notably, the NeMO Archive will be the data repository for the NIH-funded Single Cell Opioid Response in the Context of HIV (SCORCH) program as well as the BRAIN Initiative Cell Atlas Network (BICAN), the successor to the BICCN.

In the future, we hope to incorporate cell-level annotations and metadata, most importantly annotations of cell types. Currently, the NeMO Archive hosts fully-analyzed data from selected BICCN publications (12–14), including information about provenance that is consistent with the recently proposed minSCe standards (50). We will continue to ingest these data as the analyses mature. As annotated data become more prevalent, we anticipate enabling users to search the Archive for specific cell types of interest. However, these initial efforts by the BICCN are unlikely to erase the broader issue that there is no widely agreed upon catalog of brain cell types and their molecular markers. Consequently, cell type annotations are likely to change over time as new information becomes available. Also, there is no widely accepted convention for naming brain cell types, so similar cell types may be named differently across studies. Ongoing efforts from the BICCN and other groups aim to develop standards for how to define cell types for the brain based on single-cell data, extending existing standards in that area (51). The NeMO Archive is committed to building an infrastructure to support these efforts.

A related challenge is the need to integrate transcriptomic and epigenomic cell types with functional and morphological data. Despite the relatively low throughput of Patch-seq and related techniques, BICCN researchers have generated multi-modal data from thousands of neurons. In addition to the NeMO Archive, the BRAIN Initiative is supporting archives for imaging-based data (https://www.brainimagelibrary.org) and physiological data (https://dandiarchive.org/), and the Brain Cell Data Center (biccn.org) is tasked with multi-modal integration. We are working closely with these teams to ensure that users can find all the data from each cell and to support the development of multi-modal cell type models.

Finally, we are excited to support the integration of BICCN resources with efforts in the broader research community to further annotate cell types and reveal their dynamic changes across conditions. Ongoing efforts include ingestion of single-cell genomic data related to neurological disorders and the development of partnerships with other consortia and researchers developing data of these types.

## DATA AVAILABILITY

Data resources described in this manuscript are available in the NeMO Archive (nemoarchive.org) and NeMO Analytics (nemoanalytics.org).

## ACKNOWLEDGEMENTS

We thank members of the BICCN Infrastructure and Analysis Working Groups for their contributions to the development of metadata standards and data processing pipelines; all of the BICCN data submitters; Jeff Shao and Erik Anderson for IT support; and Theresa Hodges and Jain Aluvathingal for data management support.

## References Cited

1. Mukamel, E.A. and Ngai, J. (2019) Perspectives on defining cell types in the brain. Curr. Opin. Neurobiol., 56, 61–68.

2. Zeng, H. and Sanes, J.R. (2017) Neuronal cell-type classification: challenges, opportunities and the path forward. Nat. Rev. Neurosci., 18, 530–546.

3. Macosko, E.Z., Basu, A., Satija, R., Nemesh, J., Shekhar, K., Goldman, M., Tirosh, I., Bialas, A.R., Kamitaki, N., Martersteck, E.M., et al. (2015) Highly parallel genomewide expression profiling of individual cells using nanoliter droplets. Cell, 161, 1202–1214.

4. Tasic, B., Menon, V., Nguyen, T.N., Kim, T.K., Jarsky, T., Yao, Z., Levi, B., Gray, L.T., Sorensen, S.A., Dolbeare, T., et al. (2016) Adult mouse cortical cell taxonomy revealed by single cell transcriptomics. Nat. Neurosci., 19, 335–346.

5. Zeisel, A., Manchado, A.B.M., Codeluppi, S., Lonnerberg, P., La Manno, G., Jureus, A., Marques, S., Munguba, H., He, L., Betsholtz, C., et al. (2015) Cell types in the mouse cortex and hippocampus revealed by single-cell RNA-seq. Science (80-.)., 347, 1138–1142.

6. Saunders, A., Macosko, E.Z., Wysoker, A., Goldman, M., Krienen, F.M., de Rivera, H., Bien, E., Baum, M., Bortolin, L., Wang, S., et al. (2018) Molecular Diversity and Specializations among the Cells of the Adult Mouse Brain. Cell, 174, 1015–1030.e16.

7. Moffitt, J.R., Bambah-Mukku, D., Eichhorn, S.W., Vaughn, E., Shekhar, K., Perez, J.D., Rubinstein, N.D., Hao, J., Regev, A., Dulac, C., et al. (2018) Molecular, spatial, and functional single-cell profiling of the hypothalamic preoptic region. Science, 362.

8. Cadwell, C.R., Palasantza, A., Jiang, X., Berens, P., Deng, Q., Yilmaz, M., Reimer, J., Shen, S., Bethge, M., Tolias, K.F., et al. (2016) Electrophysiological, transcriptomic and morphologic profiling of single neurons using Patch-seq. Nat. Biotechnol., 34, 199–203.

9. Luo, C., Keown, C.L., Kurihara, L., Zhou, J., He, Y., Li, J., Castanon, R., Lucero, J., Nery, J.R., Sandoval, J.P., et al. (2017) Single-cell methylomes identify neuronal subtypes and regulatory elements in mammalian cortex. Science, 357, 600–604.

10. Lake, B.B., Chen, S., Sos, B.C., Fan, J., Kaeser, G.E., Yung, Y.C., Duong, T.E., Gao, D., Chun, J., Kharchenko, P. V., et al. (2018) Integrative single-cell analysis of transcriptional and epigenetic states in the human adult brain. Nat. Biotechnol., 36, 70–80.

11. Ecker, J.R., Geschwind, D.H., Kriegstein, A.R., Ngai, J., Osten, P., Polioudakis, D., Regev, A., Sestan, N., Wickersham, I.R. and Zeng, H. (2017) The BRAIN Initiative Cell Census Consortium: Lessons Learned toward Generating a Comprehensive Brain Cell Atlas. Neuron, 96, 542–557.

12. Callaway, E.M., Dong, H.-W., Ecker, J.R., Hawrylycz, M.J., Huang, Z.J., Lein, E.S., Ngai, J., Osten, P., Ren, B., Tolias, A.S., et al. (2021) A multimodal cell census and atlas of the mammalian primary motor cortex. Nature, 598, 86.

13. Bakken, T.E., Jorstad, N.L., Hu, Q., Lake, B.B., Tian, W., Kalmbach, B.E., Crow, M., Hodge, R.D., Krienen, F.M., Sorensen, S.A., et al. (2021) Comparative cellular analysis of motor cortex in human, marmoset and mouse. Nat. 2021 5987879, 598, 111–119.

14. Yao, Z., Liu, H., Xie, F., Fischer, S., Adkins, R.S., Aldridge, A.I., Ament, S.A., Bartlett, A., Behrens, M.M., Van den Berge, K., et al. (2021) A transcriptomic and epigenomic cell atlas of the mouse primary motor cortex. Nature, 598, 103.

15. Ziffra, R.S., Kim, C.N., Ross, J.M., Wilfert, A., Turner, T.N., Haeussler, M., Casella, A.M., Przytycki, P.F., Keough, K.C., Shin, D., et al. (2021) Single-cell epigenomics reveals mechanisms of human cortical development. Nature, 598, 205–213.

16. Bhaduri, A., Andrews, M.G., Mancia Leon, W., Jung, D., Shin, D., Allen, D., Jung, D., Schmunk, G., Haeussler, M., Salma, J., et al. (2020) Cell stress in cortical organoids impairs molecular subtype specification. Nature, 578, 142–148.

17. Wilkinson, M.D., Dumontier, M., Aalbersberg, Ij.J., Appleton, G., Axton, M., Baak, A., Blomberg, N., Boiten, J.W., da Silva Santos, L.B., Bourne, P.E., et al. (2016) Comment: The FAIR Guiding Principles for scientific data management and stewardship. Sci. Data, 3.

18. Schoch, C.L., Ciufo, S., Domrachev, M., Hotton, C.L., Kannan, S., Khovanskaya, R., Leipe, D., McVeigh, R., O’Neill, K., Robbertse, B., et al. (2020) NCBI Taxonomy: a comprehensive update on curation, resources and tools. Database (Oxford)., 2020.

19. Bandrowski, A., Brinkman, R., Brochhausen, M., Brush, M.H., Bug, B., Chibucos, M.C., Clancy, K., Courtot, M., Derom, D., Dumontier, M., et al. (2016) The Ontology for Biomedical Investigations. PLoS One, 11.

20. Ison, J., Kalaš, M., Jonassen, I., Bolser, D., Uludag, M., McWilliam, H., Malone, J., Lopez, R., Pettifer, S. and Rice, P. (2013) EDAM: an ontology of bioinformatics operations, types of data and identifiers, topics and formats. Bioinformatics, 29, 1325–1332.

21. Mungall, C.J., Torniai, C., Gkoutos, G. V., Lewis, S.E. and Haendel, M.A. (2012) Uberon, an integrative multi-species anatomy ontology. Genome Biol., 13.

22. Chard, K., D’Arcy, M., Heavner, B., Foster, I., Kesselman, C., Madduri, R., Rodriguez, A., Soiland-Reyes, S., Goble, C., Clark, K., et al. (2016) I’ll take that to go: Big data bags and minimal identifiers for exchange of large, complex datasets. Proc. - 2016 IEEE Int. Conf. Big Data, Big Data 2016, 10.1109/BIGDATA.2016.7840618.

23. Brase, J. (2009) DataCite - A global registration agency for research data. 4th Int. Conf. Coop. Promot. Inf. Resour. Sci. Technol. COINFO 2009, 10.1109/COINFO.2009.66.

24. Analysis Working Group, H.C.A., Data Sciences Platform, B.I. and Analysis Working Group, B.I.C.C.N. (2022) Optimus (Version 5.5.0) [Workflow].

25. Data Sciences Platform, B.I. and Analysis Working Group, B.I.C.C.N. (2021) Smartseq2_Single_Nucleus_Multisample (Version 1.1.0) [Workflow].

26. Data Sciences Platform, B.I. and Analysis Working Group, B.I.C.C.N. (2020) scATAC (Version 1.1.0) [Workflow].

27. Fang, R., Preissl, S., Li, Y., Hou, X., Lucero, J., Wang, X., Motamedi, A., Shiau, A.K., Zhou, X., Xie, F., et al. (2021) Comprehensive analysis of single cell ATAC-seq data with SnapATAC. Nat. Commun., 12.

28. Liu, H., Zhou, J., Tian, W., Luo, C., Bartlett, A., Aldridge, A., Lucero, J., Osteen, J.K., Nery, J.R., Chen, H., et al. (2021) DNA methylation atlas of the mouse brain at single-cell resolution. Nature, 598, 120–128.

29. Data Sciences Platform, B.I. and Analysis Working Group, B.I.C.C.N. (2021) CEMBA (Version 1.1.1) [Workflow].

30. Orvis, J., Gottfried, B., Kancherla, J., Adkins, R.S., Song, Y., Dror, A.A., Olley, D., Rose, K., Chrysostomou, E., Kelly, M.C., et al. (2021) gEAR: Gene Expression Analysis Resource portal for community-driven, multi-omic data exploration. Nat. Methods 2021 188, 18, 843–844.

31. Hertzano, R. and Mahurkar, A. (2022) Advancing discovery in hearing research via biologist-friendly access to multi-omic data. Hum. Genet., 141, 319–322.

32. Lim, N., Tesar, S., Belmadani, M., Poirier-Morency, G., Mancarci, B.O., Sicherman, J., Jacobson, M., Leong, J., Tan, P. and Pavlidis, P. (2021) Curation of over 10 000 transcriptomic studies to enable data reuse. Database (Oxford)., 2021.

33. Zheng, G.X.Y., Terry, J.M., Belgrader, P., Ryvkin, P., Bent, Z.W., Wilson, R., Ziraldo, S.B., Wheeler, T.D., McDermott, G.P., Zhu, J., et al. (2017) Massively parallel digital transcriptional profiling of single cells. Nat. Commun., 8, 14049.

34. Ramsköld, D., Luo, S., Wang, Y.C., Li, R., Deng, Q., Faridani, O.R., Daniels, G.A., Khrebtukova, I., Loring, J.F., Laurent, L.C., et al. (2012) Full-length mRNA-Seq from single-cell levels of RNA and individual circulating tumor cells. Nat. Biotechnol., 30, 777–782.

35. Stickels, R.R., Murray, E., Kumar, P., Li, J., Marshall, J.L., Di Bella, D.J., Arlotta, P., Macosko, E.Z. and Chen, F. (2021) Highly sensitive spatial transcriptomics at near-cellular resolution with Slide-seqV2. Nat. Biotechnol., 39, 313–319.

36. Buenrostro, J.D., Wu, B., Litzenburger, U.M., Ruff, D., Gonzales, M.L., Snyder, M.P., Chang, H.Y. and Greenleaf, W.J. (2015) Single-cell chromatin accessibility reveals principles of regulatory variation. Nature, 523, 486–490.

37. Luo, C., Rivkin, A., Zhou, J., Sandoval, J.P., Kurihara, L., Lucero, J., Castanon, R., Nery, J.R., Pinto-Duarte, A., Bui, B., et al. (2018) Robust single-cell DNA methylome profiling with snmC-seq2. Nat. Commun., 9.

38. Ma, S., Zhang, B., LaFave, L.M., Earl, A.S., Chiang, Z., Hu, Y., Ding, J., Brack, A., Kartha, V.K., Tay, T., et al. (2020) Chromatin Potential Identified by Shared Single-Cell Profiling of RNA and Chromatin. Cell, 183, 1103–1116.e20.

39. Chen, S., Lake, B.B. and Zhang, K. (2019) High-throughput sequencing of the transcriptome and chromatin accessibility in the same cell. Nat. Biotechnol., 37, 1452–1457.

40. Wang, Q., Ding, S.L., Li, Y., Royall, J., Feng, D., Lesnar, P., Graddis, N., Naeemi, M., Facer, B., Ho, A., et al. (2020) The Allen Mouse Brain Common Coordinate Framework: A 3D Reference Atlas. Cell, 181, 936–953.e20.

41. Kim, D., Langmead, B. and Salzberg, S.L. (2015) HISAT: a fast spliced aligner with low memory requirements. Nat. Methods, 12, 357–360.

42. Dobin, A., Davis, C.A., Schlesinger, F., Drenkow, J., Zaleski, C., Jha, S., Batut, P., Chaisson, M. and Gingeras, T.R. (2013) STAR: ultrafast universal RNA-seq aligner. Bioinformatics, 29, 15–21.

43. Wolf, F.A., Angerer, P. and Theis, F.J. (2018) SCANPY: Large-scale single-cell gene expression data analysis. Genome Biol., 19.

44. Chelaru, F., Smith, L., Goldstein, N. and Bravo, H.C. (2014) Epiviz: Interactive visual analytics for functional genomics data. Nat. Methods, 11, 938–940.

45. Li, M., Santpere, G., Imamura Kawasawa, Y., Evgrafov, O. V., Gulden, F.O., Pochareddy, S., Sunkin, S.M., Li, Z., Shin, Y., Zhu, Y., et al. (2018) Integrative functional genomic analysis of human brain development and neuropsychiatric risks. Science (80-.)., 362, eaat7615.

46. Nowakowski, T.J., Bhaduri, A., Pollen, A.A., Alvarado, B., Mostajo-Radji, M.A., Di Lullo, E., Haeussler, M., Sandoval-Espinosa, C., Liu, S.J., Velmeshev, D., et al. (2017) Spatiotemporal gene expression trajectories reveal developmental hierarchies of the human cortex. Science, 358, 1318–1323.

47. Jaffe, A.E., Straub, R.E., Shin, J.H., Tao, R., Gao, Y., Collado-Torres, L., Kam-Thong, T., Xi, H.S., Quan, J., Chen, Q., et al. (2018) Developmental and genetic regulation of the human cortex transcriptome illuminate schizophrenia pathogenesis. Nat. Neurosci., 21, 1117–1125.

48. Malaiya, S., Cortes-Gutierrez, M., Herb, B., Coffey, S., Legg, S., Cantle, J., Colantuoni, C., Carroll, J. and Ament, S. (2020) Single-nucleus RNA-seq reveals dysregulation of striatal cell identity due to Huntington’s disease mutations. bioRxiv, 10.1101/2020.07.08.192880.

49. Kalra, G., Milon, B., Casella, A.M., Herb, B.R., Humphries, E., Song, Y., Rose, K.P., Hertzano, R. and Ament, S.A. (2020) Biological insights from multi-omic analysis of 31 genomic risk loci for adult hearing difficulty. PLoS Genet., 16.

50. Füllgrabe, A., George, N., Green, M., Nejad, P., Aronow, B., Fexova, S.K., Fischer, C., Freeberg, M.A., Huerta, L., Morrison, N., et al. (2020) Guidelines for reporting singlecell RNA-seq experiments. Nat. Biotechnol., 38.

51. Miller, J.A., Gouwens, N.W., Tasic, B., Collman, F., Tj Van Velthoven, C., Bakken, T.E., Hawrylycz, M.J., Zeng, H., Lein, E.S. and Bernard, A. Common cell type nomenclature for the mammalian brain. elifesciences.org, 10.7554/eLife.59928.

